# Food collection behavior of *Apis mellifera* and *Tetragonisca angustula* bees in *Brassica napus* L. in response to different environmental covariates: Environmental covariates and foraging behavior in *Brassica napus* L.

**DOI:** 10.1101/428128

**Authors:** Simone Cristina Camargo, Regina Conceição Garcia, Newton Tavares Escocard de Oliveira, Edmar Soares de Vasconcelos, Douglas Galhardo, Sandra Mara Ströher, Alceu Maurício Hartleben, Bruno Garcia Pires, Gustavo Andrés Patiño Piñeros, David Fernando Gómez Díaz, Renato de Jesus Ribeiro, Thiago Henrique Radtke, Pedro da Rosa Santos, Vagner de Alencar Arnaut de Toledo

**Author notes:** These authors contributed equally to this work (Simone Cristina Camargo, Regina Conceição Garcia).

## Abstract

The objective was to evaluate the behavior of *Apis mellifera* and *Tetragonisca angustula* bees in pollination tests in *Brassica napus* at different times of the day, temperature and relative humidity. The experimental design was completely with eight treatments and two pollination tests, repeated in five randomized blocks during seven days of observations for two years, totaling 560 records. During the visits, the following parameters were recorded: collected resources, nectar collection site, time spent on flower, number of flowers visited in one minute, pollen load in the pollen basket and bee contact with anthers and stigma. Data were analyzed using generalized linear models. The number of Africanized and *T. angustula* bees collecting nectar increased with the passage of time throughout the day and with the decrease of relative humidity. The same was observed for nectar collection in both nectaries. The proportion of bees collecting pollen was higher in the morning hours, as well as when there was an increase in temperature for the two species of bees. Foraging behavior of *A. mellifera* in *B. napus* crop favored its pollination, indifferent of which collected floral resource as they came into contact with anthers and stigma. *T. angustula* bees performed pollination only during pollen collection. Pollination of *B. napus* was more effective in the warmer hours of the morning, when more of both species of bees carried out pollen collection. Due to its foraging behavior, *A. mellifera* had greater efficiency for pollination of *B. napus*; however, the association with T*. angustula* may potentiate the benefits generated for the crop by cross-pollination.

## Introduction

Pollination is considered an ecosystem service which provides benefits to plants and pollinators, increasing the adaptive value (fitness) of both [1]. Also, pollination is important for biodiversity conservation and food production. In agriculture, both the farmer and the beekeeper benefit from pollinating through increased productivity [2]. Estimates performed by [3] indicated that 87.5% of angiosperms are pollinated by animals and, on average, 33% of major agricultural crops depend to some degree on insect pollination.

The pollination service by bees in Brazil generates a revenue of US$ 12-14 billion [4]. Among the most important crops is the canola (*Brassica napus* L. var. oleifera), which is the third most produced oilseed in the world and has been used as biodiesel [5]. This plant has hermaphroditic flowers and it is self-compatible, and may produce fruits and seeds by both self-pollination and by cross-pollination [6]. Studies have shown that cross-pollination can increase production [7] and the economic value of canola grains [6].

The dispersal of canola pollen grains is mediated by a variety of pollinating insects, which are attracted by the aroma and color of their flowers, as well as by their nutritional resources [8]. *B. napus* flowers have an average nectar production ranging from 0.02 to 0.75 µL per flower [9] and produce on average 9 kg/ha of pollen per day [10].

To an efficient pollination process, it is necessary to the floral visitor to present specific morphological and behavioral characteristics for each type of flower [1]. This is the case of *Apis mellifera* honeybees that have been identified as effective canola pollinators in different parts of the world [7;11;12].

In spite of the recognized efficiency of *A. mellifera* as pollinator, the identification of alternative pollinators is also important because, in addition to encouraging their conservation, it ensures the continuity of pollination services in case of decrease or disappearance of the main pollinator [13]. In canola cultivation, species of *Halictus* sp. [8], *Bombus lapidarius* [14], and *Colletes lacunatus* [11] bees were identified as alternative pollinators. In Brazil, the stingless bee species *Trigona spinipes* was identified with a potential pollinator for canola [15].

Considering the importance of native bees in pollination of agricultural crops [16], especially of *Tetragonisca angustula* for the western region of Paraná, as well as the variation in bees’ foraging behavior [15], the objective of this study was to evaluate the time effects on pollination tests and environmental covariates (relative humidity and temperature) at different times of the day on the behavior of *A. mellifera* and *T. angustula* bees during foraging in flowers of canola (*B. napus*).

## Material and methods

The experiment was developed at the Experimental Station Professor Alcibíades Luiz Orlando, of the State University of the Western Paraná - UNIOESTE, in the county of Entre Rios do Oeste, State of Parana, Brazil (24º40′51″S and 54º16′56″W) and 400 meters above sea level.

The evaluated cultivar was Hyola 76 of long cycle, that lasts from 120 to 164 days from the emergence of the seedling until maturation [17]. The seeds were sown on 05/31/2015 and 05/05/2016, under a no-tillage system, with soybeans as their predecessor crop.

In the first year, flowering occurred on 05/08/2015 and extending until 12/09/2015, in the second year, the beginning of flowering was 12/07/2016 remaining until 08/27/2016. In the two years, the meteorological data (temperature and relative air humidity) were collected inside the pollination cages using a thermohydrometer.

The experiment was set up in a completely randomized block design, with eight treatments repeated 14 times in each block, totaling 560 records. The blocks were composed of five sets containing two pollination cages each. The treatments were comprised of the combination of four times of day (9:00 a.m., 11:30 a.m. 13:30 p.m. and 15:30 p.m.) and two pollination tests: area covered with a pollination cage with a colony of *A. mellifera* inside and area covered with pollinating cage with *T. angustula*.

The behavior of the foragers honeybees was analyzed by direct observations of six bees at each time hour during their visits to the flowers. For each visit, the following variables were recorded: time spent on flower, number of flowers visited in one minute, type of floral resource collected (nectar or pollen), specific nectar collection site (lateral or median nectary), pollen load in the pollen basket, and if there was contact with the anthers and stigma.

A nuc hive with five frames was used for *A. mellifera*, three brood combs and two with storage food in each nuc, while for *T. angustula* standard hives were used with a nest and two honey supers, with internal measurements of 20 cm × 20 cm × 8 cm. The colonies were inserted in the crop when it was 10% flowering on 08/08/2015 and 07/18/2016, and during the whole flowering period the colonies individually received drinking water and sugar syrup in the concentration of 25 mol/L as food supplement.

The pollination cages were made following the model proposed by [20], with a 2 × 2 mm mesh nylon screen, supported by ¾ inch PVC tubes, 4 m wide, 6 m long, and 2 m high at the top, totaling an area of 24 m^2^ (Fig 1). They were assembled before flowering, on the dates of 07/31/2015 in the first year and 05/07/2016 for the second year, remaining in the crop until the end of the flowering period.

**Fig 1.**
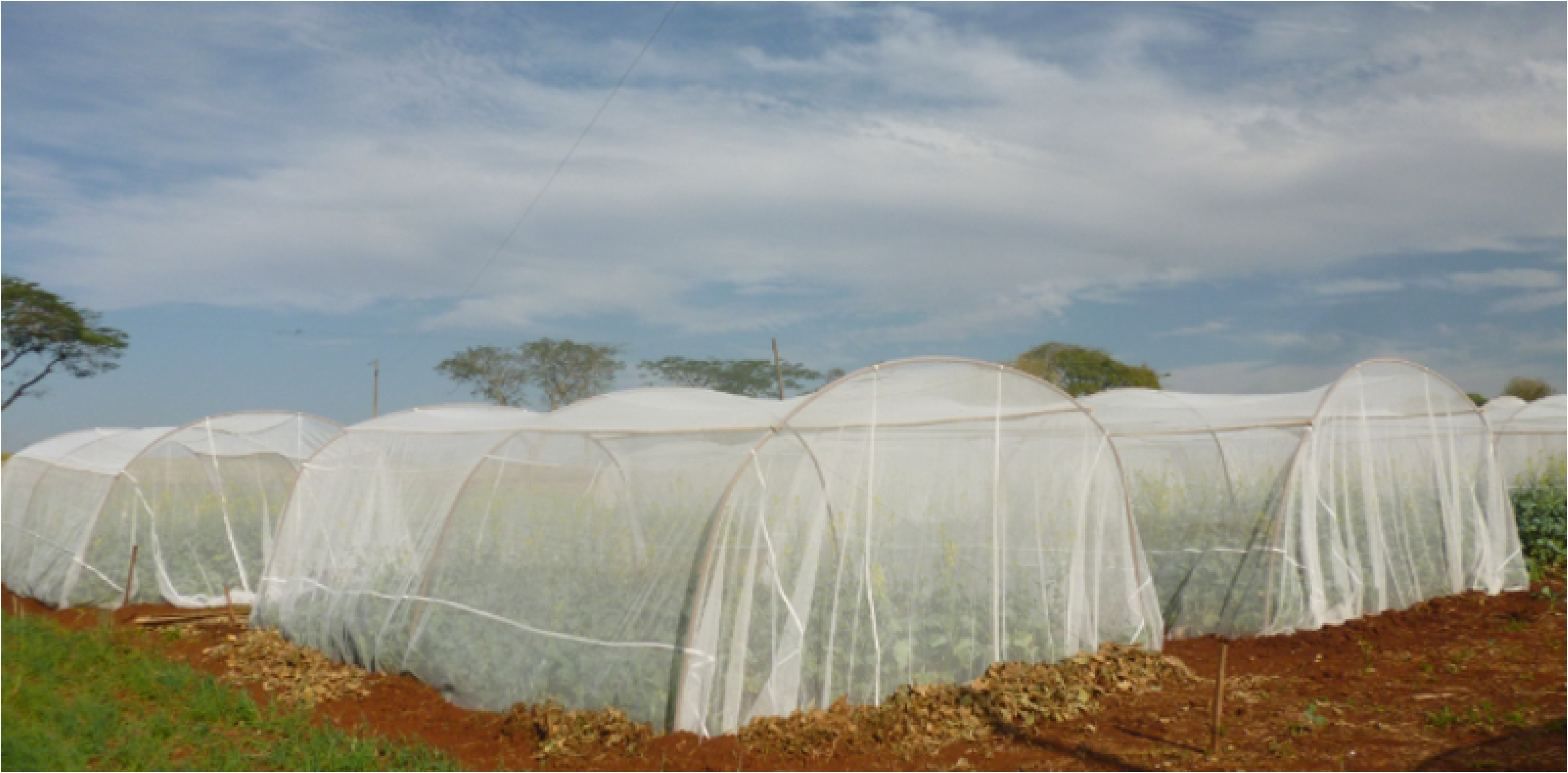
Pollination cages installed in the canola crop.

For the statistical procedures, generalized linear models (GLM), with binomial error structure and logit link function expressed by g(µ) = ln(µ/1-µ), were used to verify the effects of time (hours) and pollination test on the following variables: number of bees collecting nectar, number of bees collecting pollen, number of bees collecting nectar and pollen, number of bees collecting in the median nectaries, number of bees collecting in lateral nectaries, number of bees collecting in the median and lateral nectaries, number of bees with pollen in the pollen basket and number of bees that came in contact with anthers and stigma. The number of bees that came into contact with anthers only was considered as an error, with normal distribution g(μ) = μ. Mean time spent on flower and the average number of flowers visited by each bee in one minute were adjusted to the Poisson distribution, with logarithmic function g(μ) = ln(μ).

The parameters of the model were estimated using the maximum likelihood method by maximizing the log-likelihood function, using generalized estimation equations (GEE). In the GEE adjustment, the dependence between observations measured within time (hours), which characterized repeated measures in the experimental units (pollination cages), was considered. The responses obtained between the experimental units of different blocks were considered independent.

*Deviance* analysis was used to adjust the GLM from a model represented by systematic portion:

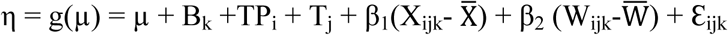

in which:

µ is the effect of the general average;
B_k_ is the block effect (k = 1, 2, 3, 4, and 5);
TP_i_ is the effect of the pollination test (i = 1 and 2);
T_j_ is the effect of time (j = 1, 2, 3, and 4);
β_1_ is the regression coefficient of g(µ) about X;
X_ijk_ is the observation of the covariable “temperature” in each plot;
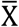 is the general mean temperature;
β_2_ is the regression coefficient of g (μ) about W;
W_ijk_ is the observation of the covariable “relative humidity” in each plot;
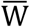 is the general average relative humidity; and
Ɛ_ijk_ is the random error of the plots.

In the occurrence of a higher probability of significance for a given factor, it was removed from the statistical model and the analysis was reprocessed. The fit quality of the models to the observed data was based on the logarithm of the maximum likelihood function (*LogLik*). The dispersion parameters were adjusted for superdispersion, correcting the standard errors using model *quasibinomial*, *quasipoisson*, and normal for the respective variables mentioned above.

The block effect, pollination test, time, temperature, and relative humidity on the variables were verified by Type 3 GEE analysis. When significance occurred (p ≤ 0.05), the pollination, time, temperature, and relative humidity effects on the variables were evaluated through logistic regression models.

A level of significance of 0.05 was adopted in all hypothesis tests. Statistical analyses were performed using the software R (R Core Development Team, 2018) [18].

## Results

*A. mellifera* honeybees collecting only nectar were more frequent (67.68%) than those that collected only pollen (11.75%) or both resources (20.57%). A higher frequency of nectar collectors was also observed in *T. angustula* bees (74.31%), when compared to that of pollen collectors and that of both resources (24.81 and 0.88%, respectively). The percentage of bees carrying pollen load in the pollen basket was 35.53% in *A. mellifera* and 23.93% in *T. angustula*.

The collection of nectar by *A. mellifera* honeybees was performed mainly in the median and lateral nectaries in the same visit (84.14%), followed by lateral nectaries (14.60%) and median nectaries (1.26%). The Africanized honeybees perched on the corolla and, with the proboscis extended between the openings of the petals, they sucked the nectar present in the lateral nectaries. Then they walked on the flower for collection in the median nectaries, touching the reproductive organs (anthers and stigma) of *B. napus* flowers.

*T. angustula* collected nectar mainly in the median nectaries (46.02%), followed by the median and lateral (45.52%), and to a lesser extent in the lateral (8.46%). In order to reach the median nectaries, *T. angustula* bees lay on the side of the flower, between the sepals and, approaching the base of the short stamens on the side, with the extended proboscis were able to reach the nectar present in the lateral nectaries, without coming into contact with the reproductive organs of flowers.

During the visits, 100% of *A. mellifera* foragers observed came into contact with anthers and stigma, while 68.11% of *T. angustula* came in contact with the anthers only and 31.89% had contact with the anthers or the anthers and stigma. Africanized honeybees averaged 13.20 ± 1.74 flowers in one minute, and remained on average 4.10 ± 0.67 seconds per flower. *T. angustula* bees averaged 4.30 ± 1.59 flowers in one minute and the mean time spent on *B. napus* flowers was 12.10 ± 3.94 seconds.

The quality of fit of GLM to behavioral data during the foraging of *A. mellifera* and *T. angustula* bees on *B. napus* was evaluated by the “scaled deviance” statistic (Table 1).

**Table 1.** Results of the “scaled deviance” values* to measure the quality of fit of the generalized linear models.

| Variable | Degrees<br>of<br>freedom | “Deviance”<br>Value | Maximum<br>log likelihood |
| --- | --- | --- | --- |
| Bees collecting nectar | 556 | 556.0 | -567.8 |
| Bees collecting nectar in the median nectaries | 556 | 556.0 | -666.3 |
| Bees collecting nectar in the lateral nectaries | 556 | 556.0 | -382.1 |
| Bees collecting nectar in the median and lateral nectaries | 556 | 556.0 | -528.3 |
| Bees collecting pollen | 555 | 555.0 | -640.2 |
| Bees collecting nectar and pollen | 557 | 557.0 | -622.7 |
| Pollen load in the pollen basket | 555 | 555.0 | -591.3 |
| Time spent on flower | 555 | 555.0 | 7248.7 |
| Number of flowers visited in one minute by the bees | 556 | 556.0 | 14562.5 |
| Bees that came in contact with anthers and stigma | 557 | 557.0 | -679.3 |
| Bees that came in contact with anthers | 557 | 557.0 | -3068.6 |

\* Scaled deviance degrees of freedom were equal to 1.0 for all variables, due to the use of the "scaled deviance" command, which corrects the standard errors of the parameters.

From the summary of the Type 3 statistical analysis, analyzing the probability values (*P*) related to the sources of variation included in the statistical model for the proportion of bees *A. mellifera* and *T. angustula* collecting nectar, there were time (*P* = 0.0010) and relative humidity (*P* < 0.0001) effects. There were effects of temperature (*P* = 0.0006 and *P* < 0.0001) and relative humidity (*P* = 0.0032 and *P* < 0.0001) on the proportion of *A. mellifera* and *T. angustula* that collected nectar in the median and lateral nectaries, respectively. For the two species of bees that collected in the median and lateral nectaries in the same visit, there were effects of time (*P* = 0.0010) and relative humidity (*P* < 0.0001).

An effect was observed for time (*P* < 0.0001), temperature (*P* = 0.0006), and relative humidity (*P* < 0.0001) on the proportion of Africanized and *T. angustula* bees collecting pollen. There was a relative humidity effect (*P* < 0.0001) in the proportion of *A. mellifera* and *T. angustula* collecting nectar and pollen. There was an effect of time (*P* = 0.0001 and *P* = 0.0161), temperature (*P* < 0.0001 and *P ≤* 0.0001), and relative humidity (*P* < 0.0001 and *P* = 0.0004) for the proportion of *A. mellifera* and *T. angustula* with pollen load in the pollen basket and for the time spent on flower, respectively. The number of flowers visited in one minute by *A. mellifera* and *T. angustula* foragers was affected by the time and relative humidity (*P* = 0.0130 and *P* < 0.0001, respectively). The proportions of *A. mellifera* and *T. angustula* foragers that came in contact with the anthers and stigma or only with the anthers were influenced by the relative humidity (*P* < 0.0001 and *P* = 0.0002, respectively).

Significant estimates were found for the parameters of the models (*P* < 0.05), with the exception of the intercepts in the adjusted models for the variables: proportion of *A. mellifera* and *T. angustula* collectors in the median and lateral nectaries and proportion of bees that came into contact with anthers only (Table 2).

**Table 2.**
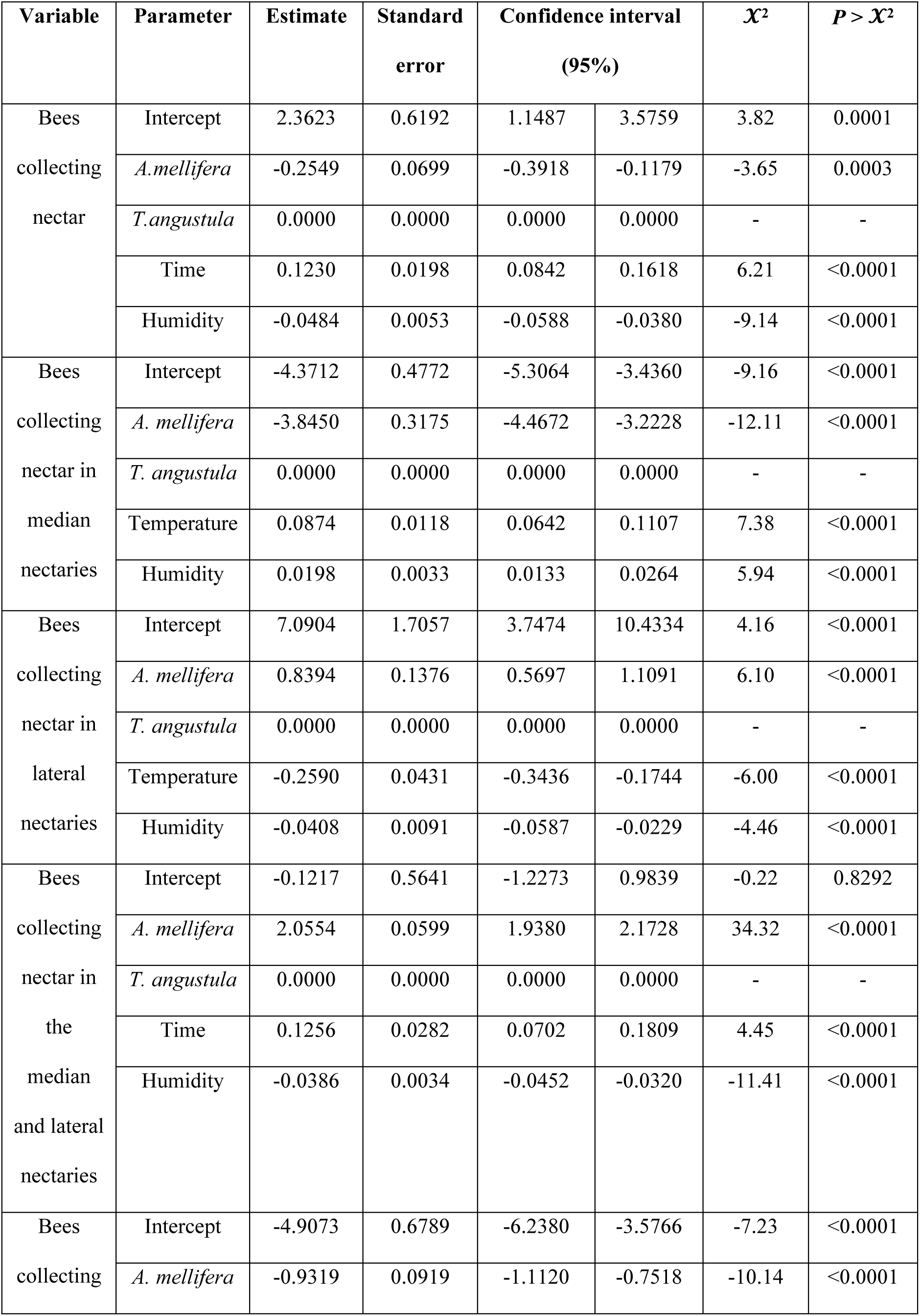
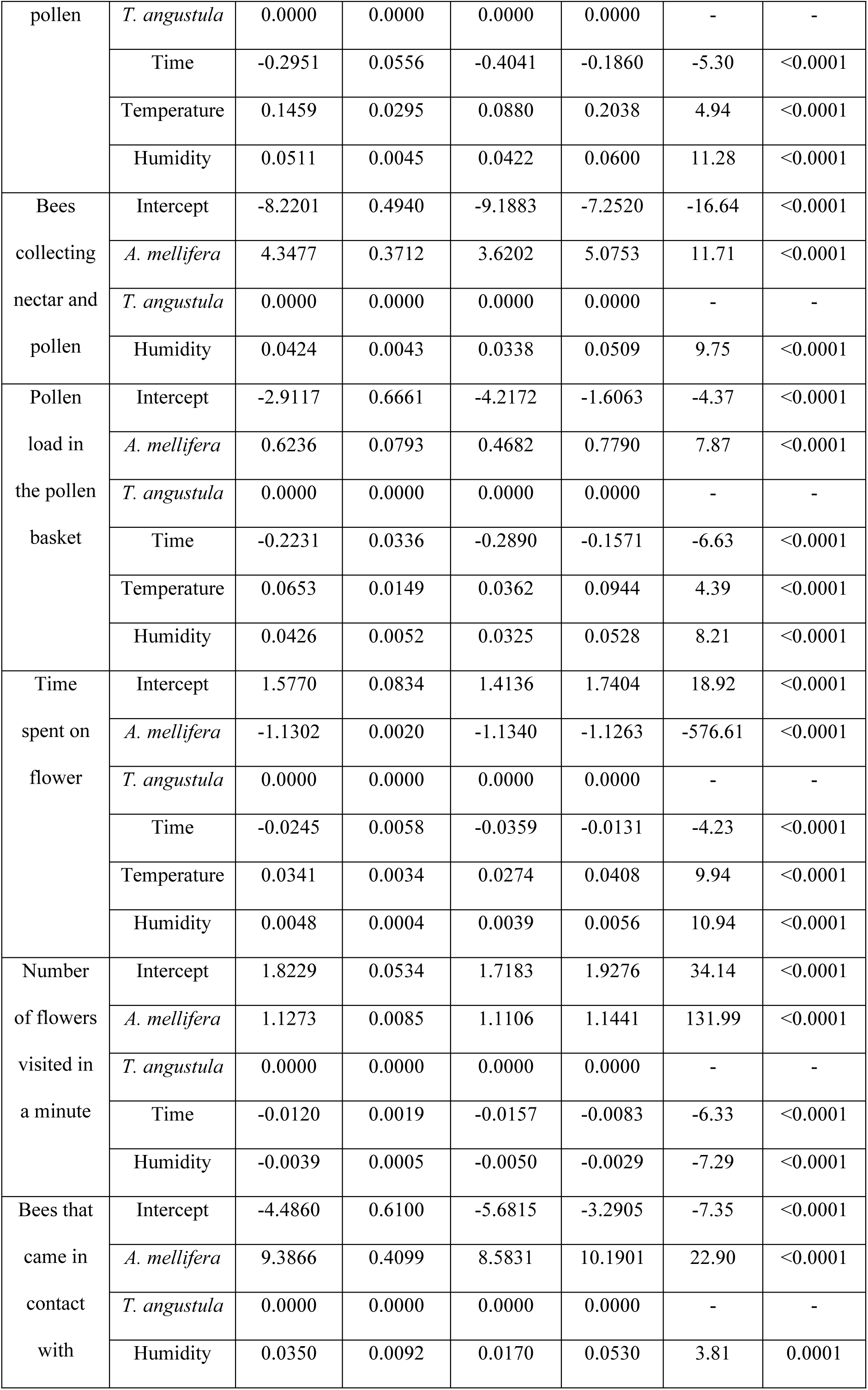
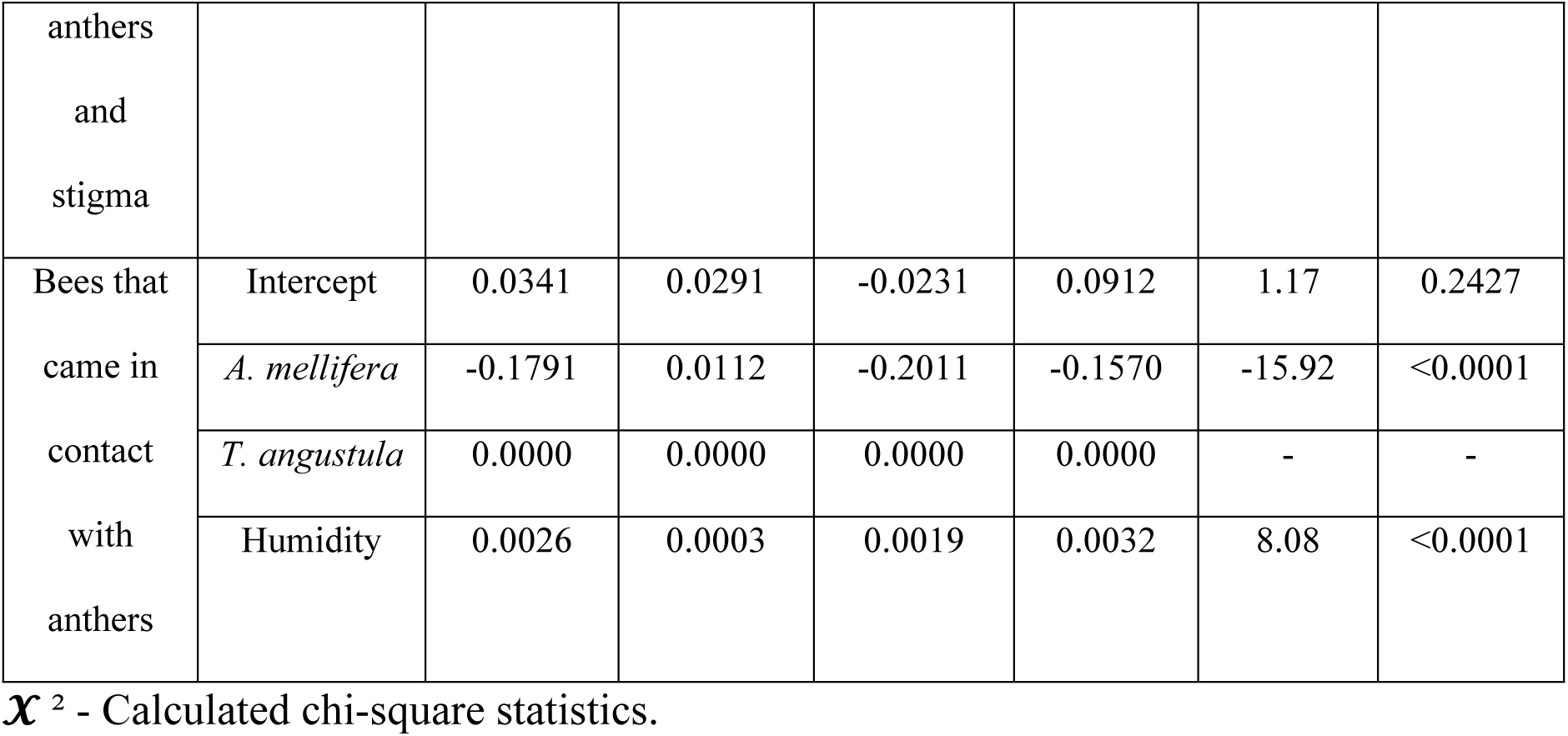
Analysis of the estimates of the parameters of the regression models and respective descriptive levels.

From the regression equations estimated for each species of bee, it was observed that the exclusive behavior of collecting nectar or pollen was more frequent in *T. angustula*. These bees also performed more nectar collection from median nectaries. The time spent on flower was greater than that of *A. mellifera*. The behavior of nectar and pollen collection, including collection in the lateral nectaries and in both nectaries (median and lateral), as well as the presence of pollen in the pollen basket were more frequent in *A. mellifera* foragers. During their visits to the flowers these foragers came in contact with the anthers and stigma, visiting a larger number of flowers compared to *T. angustula*.

The estimated regression equations for both species showed that the average number of Africanized and *T. angustula* bees collecting nectar increased with the passage of hours during the day, and with the decrease of relative humidity (Fig 2). The number of *A. mellifera* and *T. angustula* bees collecting nectar only in the median nectaries was higher when the relative humidity and temperature increased (Fig 3).

**Fig 2.**
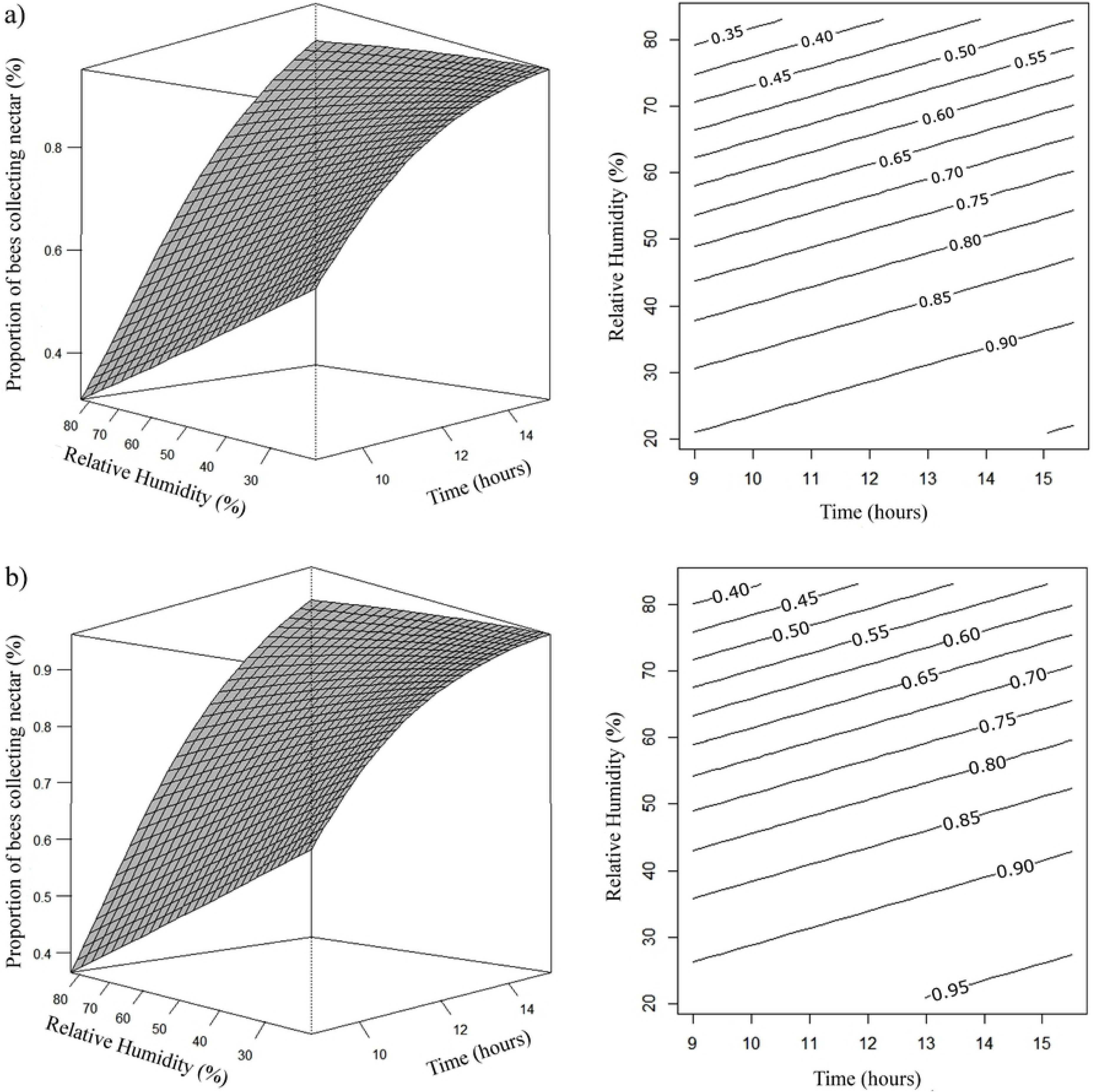
Response surface (left side) and contour (right side) graphics. Effects of daytime (x) and relative humidity (rh) on the proportion (μ) of *A. mellifera* (a) and *T. angustula* (b) bees collecting nectar. (a) µ= e^2,1074+0,1230x–0,0484rh^/1 + e^2,1155+0,1230x−0,0484rh^; (b) µ= e^2,3623+0,1230x–0,0484rh^/1 + e^2,3698+0,1230x–0,0484rh^.

**Fig 3.**
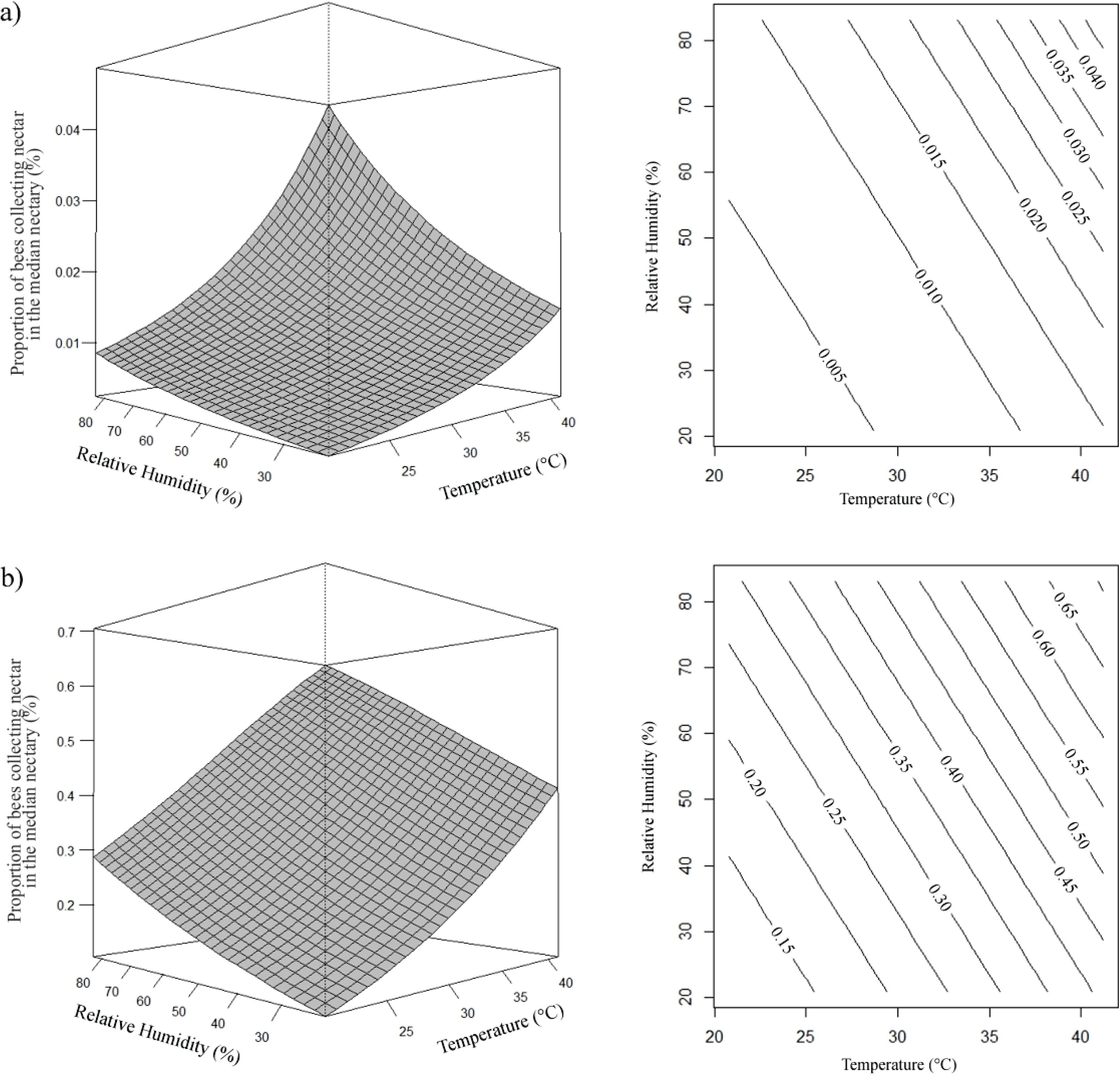
Response surface (left side) and contour (right side) graphics. Effects of temperature (temp) and relative humidity (rh) on the proportion (μ) of *A. mellifera* (a) and *T. angustula* (b) bees collecting nectar in the median nectaries. (a) µ= e^−8,2162+0,0874temp+0,0198rh^/1 + e^−8,2162+0,0874temp+0,0198rh^; (b) µ= e^−4,3712+0,0874temp+0,0198rh^/1 + e^−4,3712+0,0874temp+0,0198rh^.

The opposite was observed in the estimate of the number of bees collecting nectar only in the lateral nectaries, as the relative humidity and temperature decreased the proportion of bees collecting nectar in the lateral nectaries was higher (Fig 4). It was verified that estimated proportion of *A. mellifera* and *T. angustula* bees collecting nectar in the median and lateral nectaries, increased with decreasing relative humidity and as the time passed, being higher in the evening hours (Fig 5).

**Fig 4.**
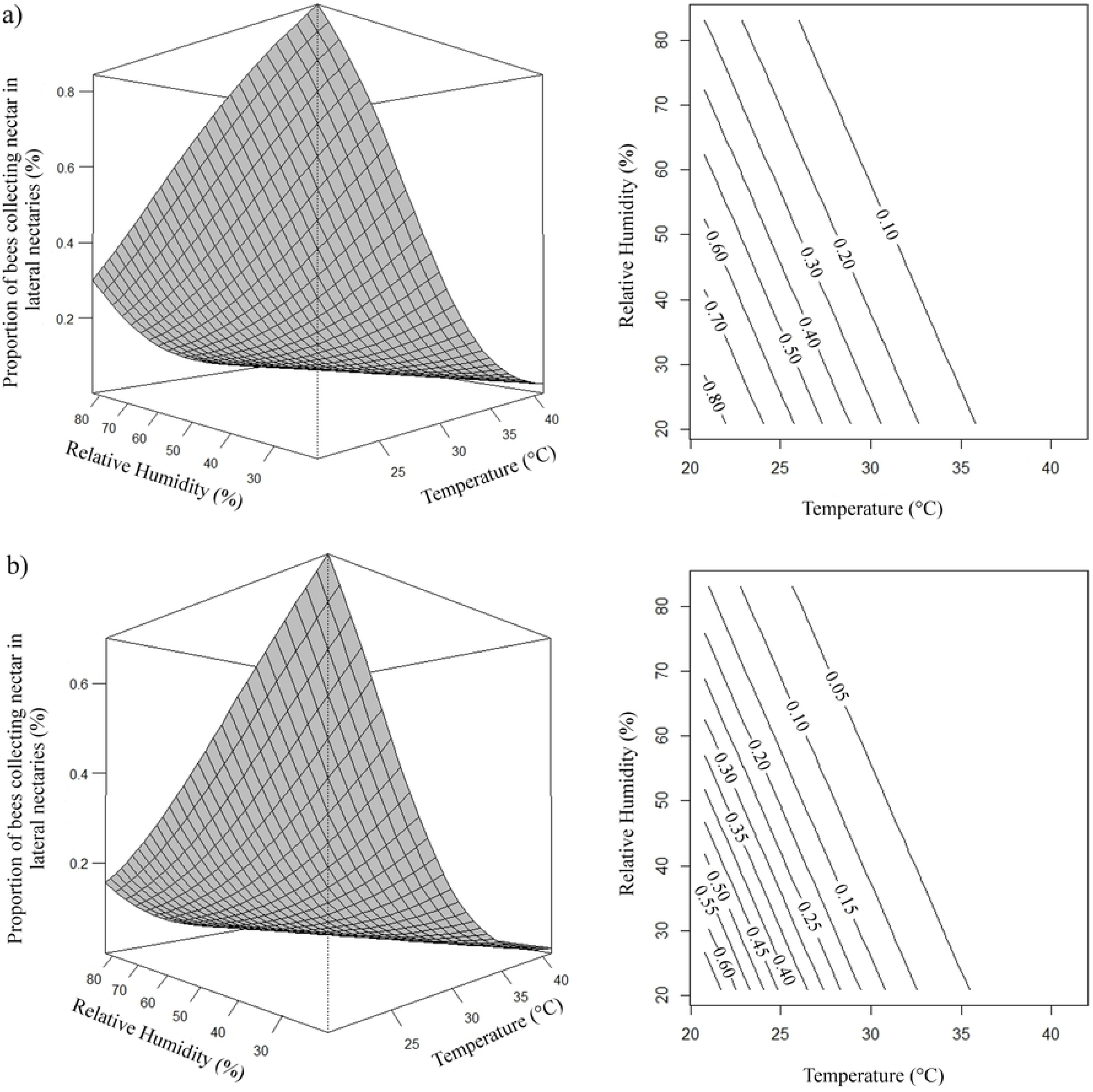
Response surface (left side) and contour (right side) graphics. Effects of temperature (temp) and relative humidity (rh) on the proportion (μ) of *A. mellifera* (a) and *T. angustula* (b) bees collecting nectar in lateral nectaries. (a) µ= e^7,9298-0,2590temp-0,0408rh^/1 + e^7,9298-0,2590temp-0,0408rh^; (b) µ= e^7,0904-0,2590temp-0,0408rh^/1 + e^7,0904-0,2590temp-0,0408rh^.

**Fig 5.**
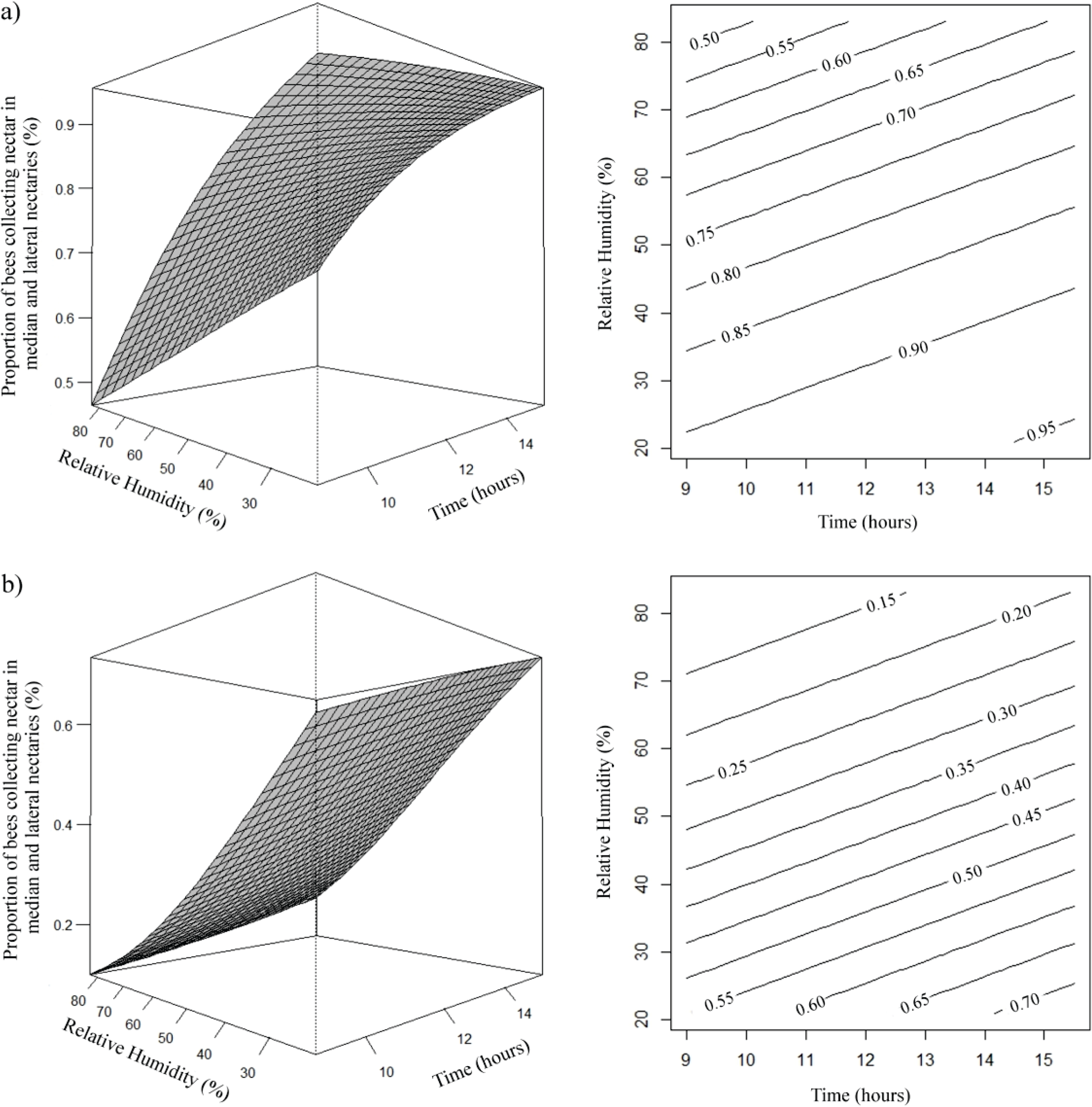
Response surface (left side) and contour (right side) graphics. Effects of daytime (x) and relative humidity (rh) on the proportion (μ) of *A. mellifera* (a) and *T. angustula* (b) bees collecting nectar in the median nectaries and lateral nectaries. (a) µ= e^1,9337+0,1256x–0,0386rh^/1 + e^1,9337+0,1256x–0,0386rh^; (b) µ= e^−0,1217+0,1256x-0,0386rh^/1 + e^−0,1217+0,1256x-0,0386rh^.

For the proportion of bees collecting pollen, it was shown that the estimate was higher with the increase of temperature and relative humidity and in the morning hours, decreasing with the hours of the day for both species (Fig 6). It was observed that estimation of bees collecting nectar and pollen increased with increasing relative humidity, especially above 65%, for both *A. mellifera* and *T. angustula* (Fig 7).

**Fig 6.**
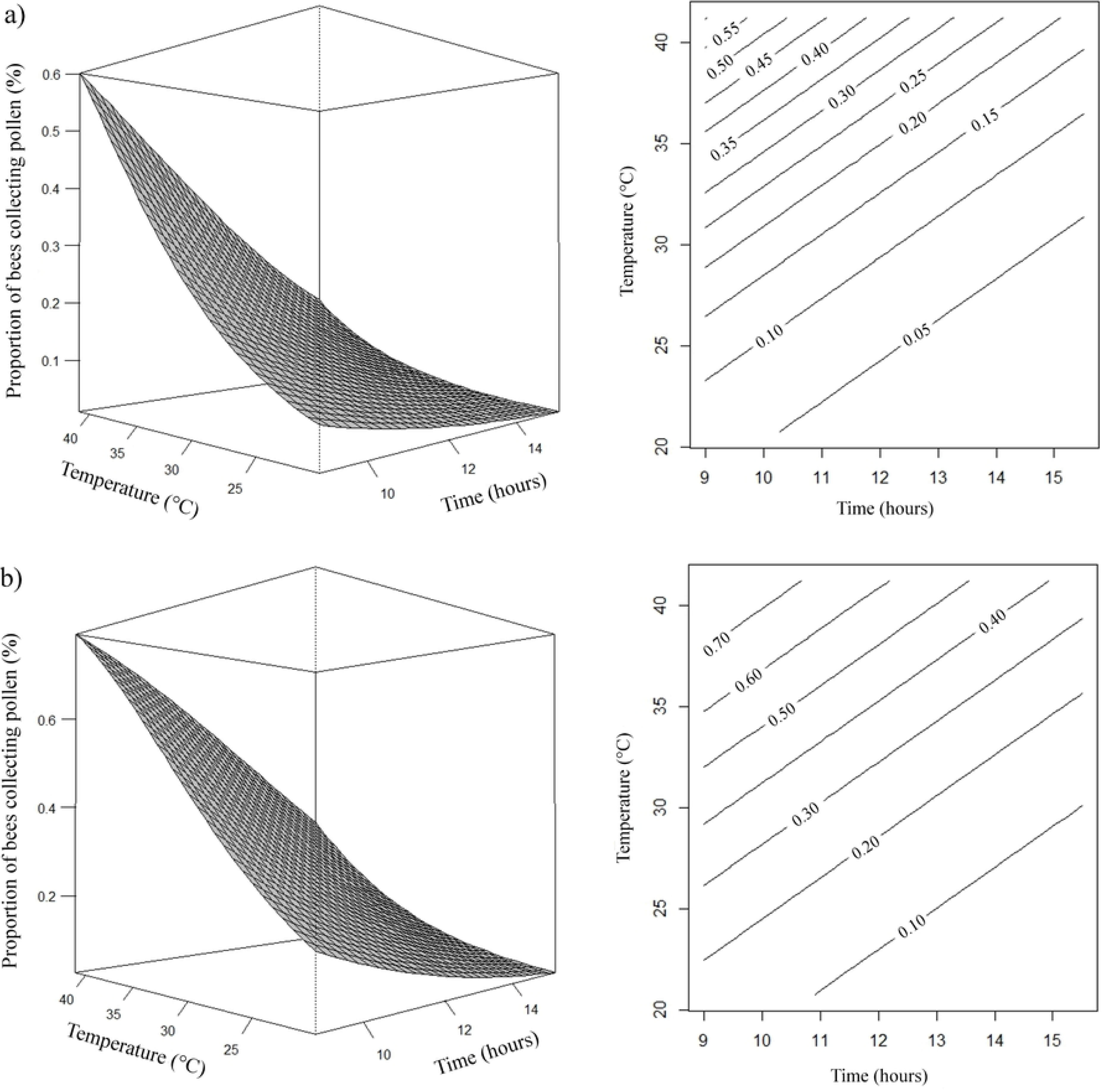
Response surface (left side) and contour (right side) graphics. Effects of daytime (x) and temperature (temp) on the proportion (μ) of *A. mellifera* (a) and *T. angustula* (b) bees pollen collectors. (a) µ= e^−2,9449–0,2951x+0,1459temp^/1 + e^−2,9449– 0,2951x+0,1459temp^; (b) µ= e^−2,0130–0,2951x+0,1459temp^/1 + e^−2,0130–0,2951x+0,1459temp^; Cut-off point of the average relative humidity at 56.64%.

**Fig 7.**
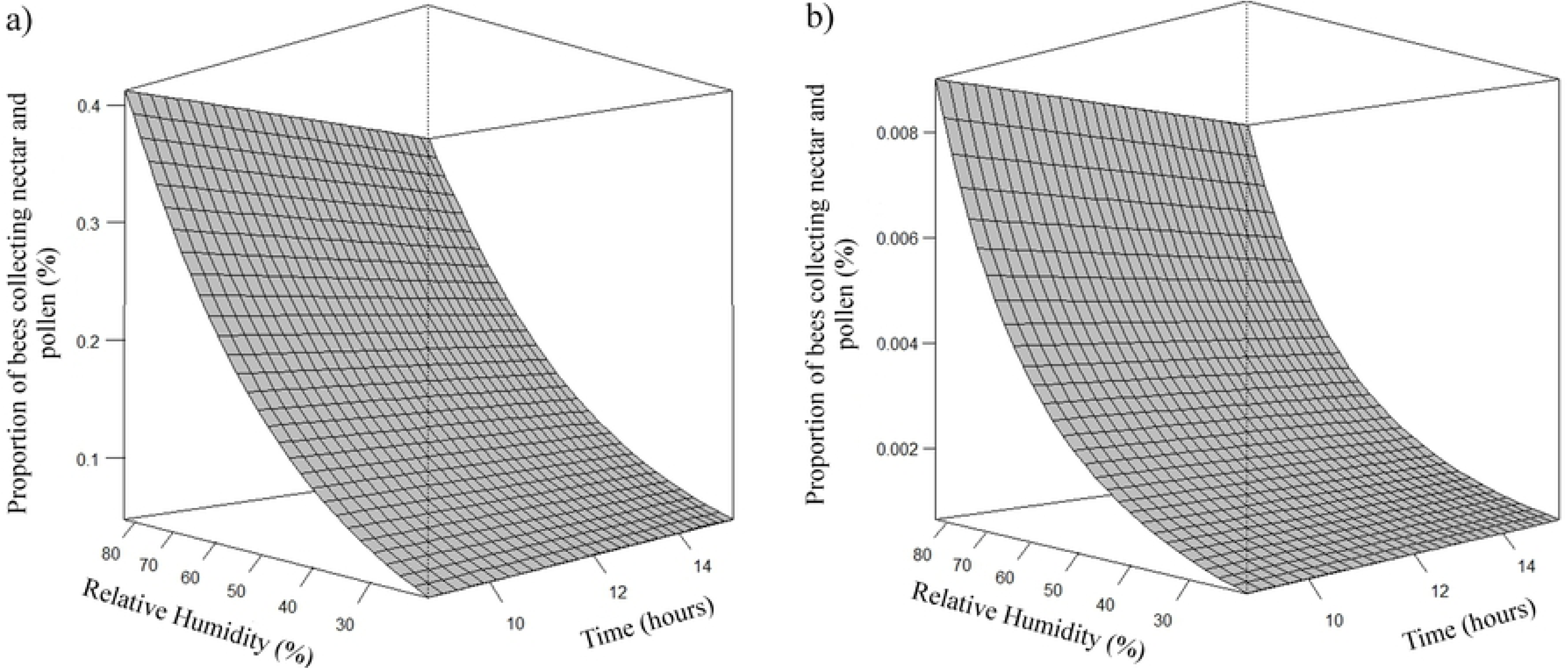
Response surface graphic. Effect of relative humidity (rh) on the proportion (μ) of *A. mellifera* (a) and *T. angustula* (b) bees’ collectors of nectar and pollen. (a) µ= e-^3,8724+0,0424rh^/1+ e^−3,8724+0,0424rh^; (b) µ= e^−8,2201+0,0424rh^/1 + e^−8,2201+0,0424rh^.

The estimate of the average proportion of bees carrying pollen load in the pollen basket was higher when there was an increase in temperature and relative humidity, decreasing as the hours of the day passed, for both species of bees (Fig 8). For the time spent on flower, the estimates were higher when relative humidity and higher temperatures occurred, mainly above 32ºC, along with the morning period (up to 10:30 a.m.), decreased with the passage of the day, for Africanized and *T. angustula* bees (Fig 9).

**Fig 8.**
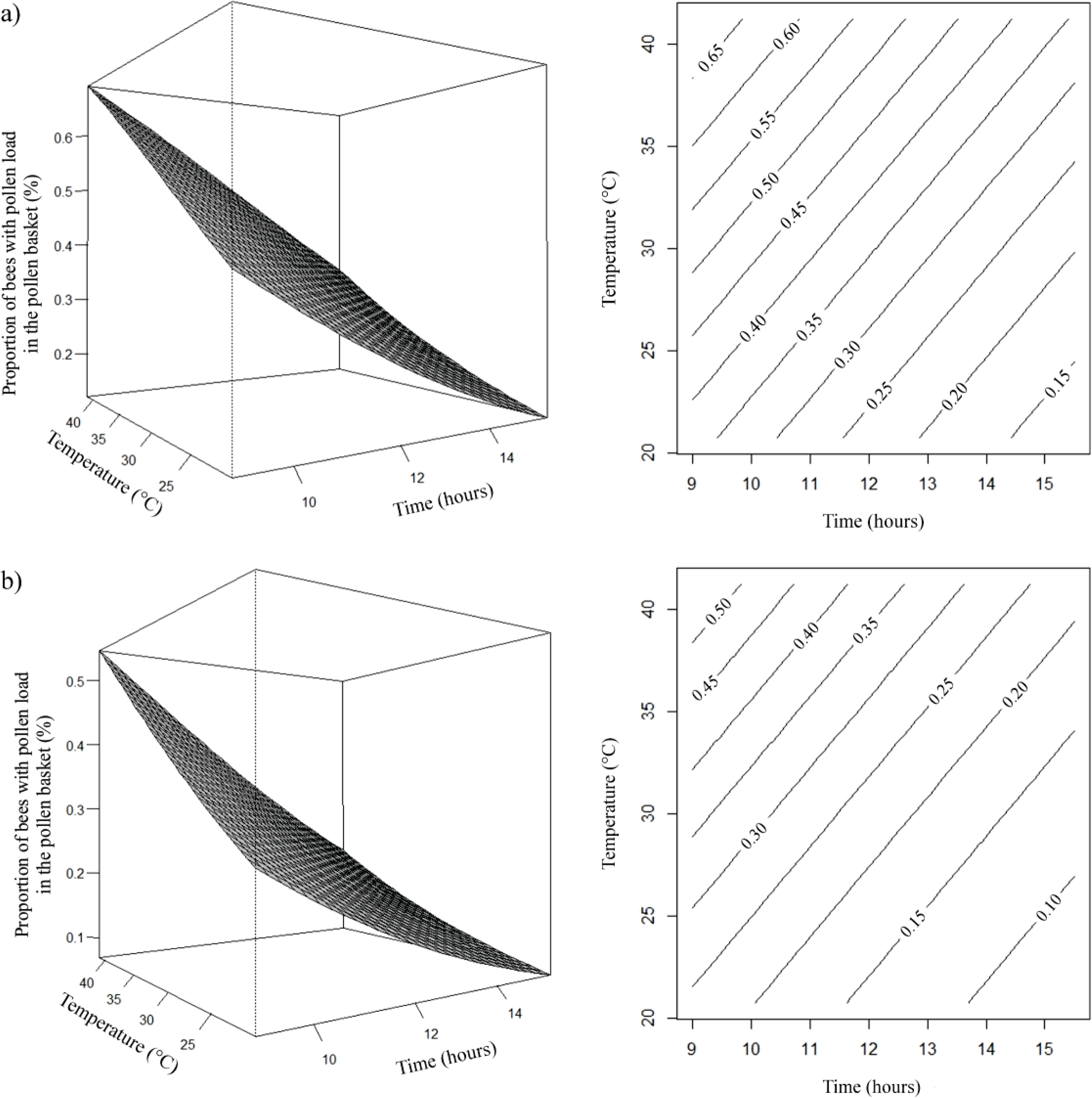
Response surface (left side) and contour (right side) graphics. Effects of daytime (x) and temperature (temp) on the proportion (μ) of *A. mellifera* (a) and *T. angustula* (b) bees carrying pollen load in the pollen basket. (a) µ= µ= e^0,1248-0,2231x+0,0653temp^/1 + e^0,1248-0,2231x+0,0653temp^; (b) µ = e^−0,4988-0,2231x+0,0653temp^/1 + e^−0,4988-0,2231x+0,0653temp^; Cut-off point of the average relative humidity at 56.64%.

**Fig 9.**
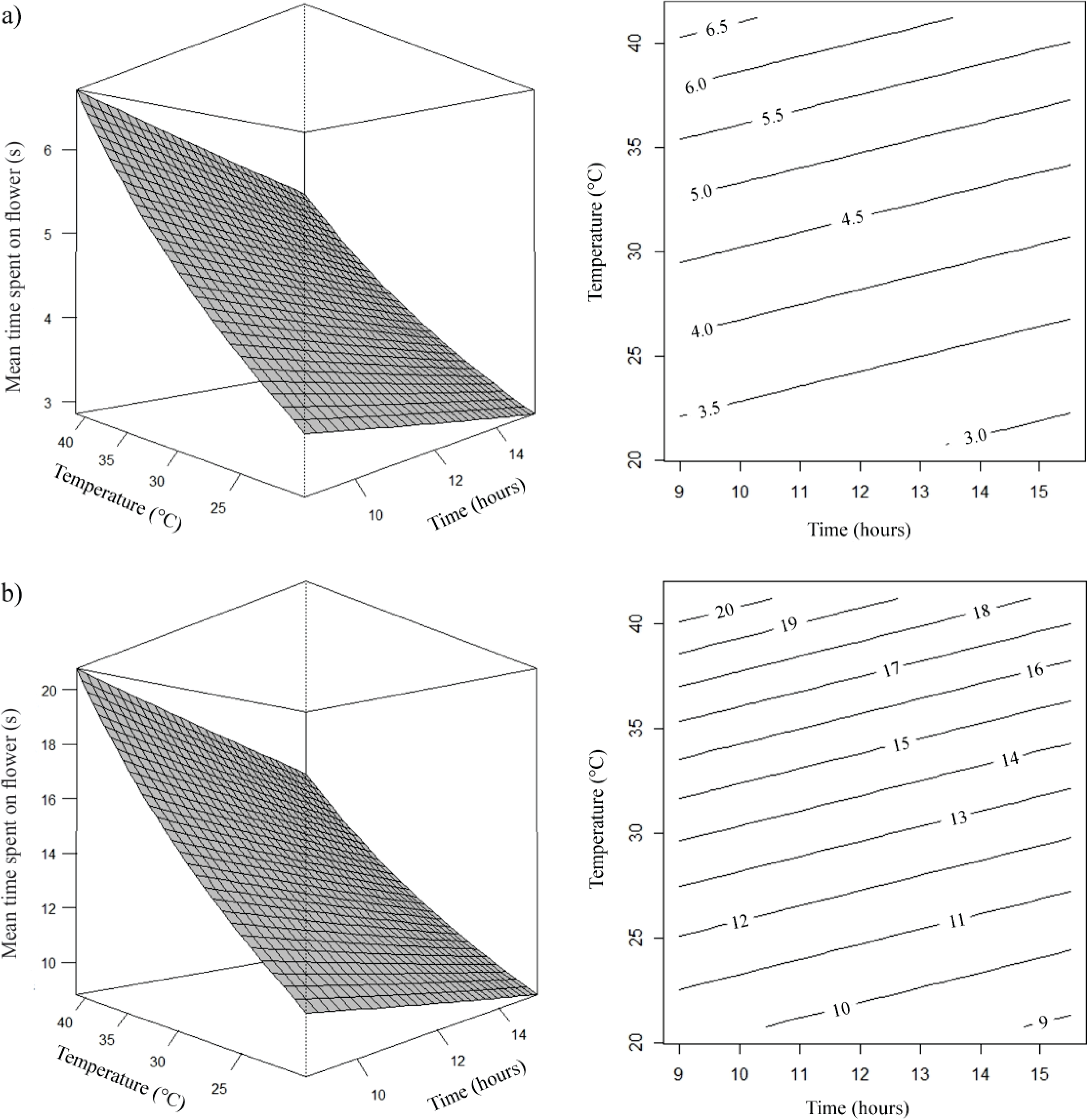
Response surface (left side) and contour (right side) graphics. Effects of daytime (x) and temperature (temp) on the average time (μ) of *A. mellifera* (a) and *T angustula* (b) bees on the bloom. (a) µ= e^0,7187-0,0245x+0,0341temp^; (b) µ= e^1,8489-0,0245x+0,0341temp^; Cut-off point of the average relative humidity at 56.64%.

For both species the number of flowers visited in one minute was higher when the relative humidity was lower and in the morning period, decreased with the passage of the day (Fig 10).

**Fig 10.**
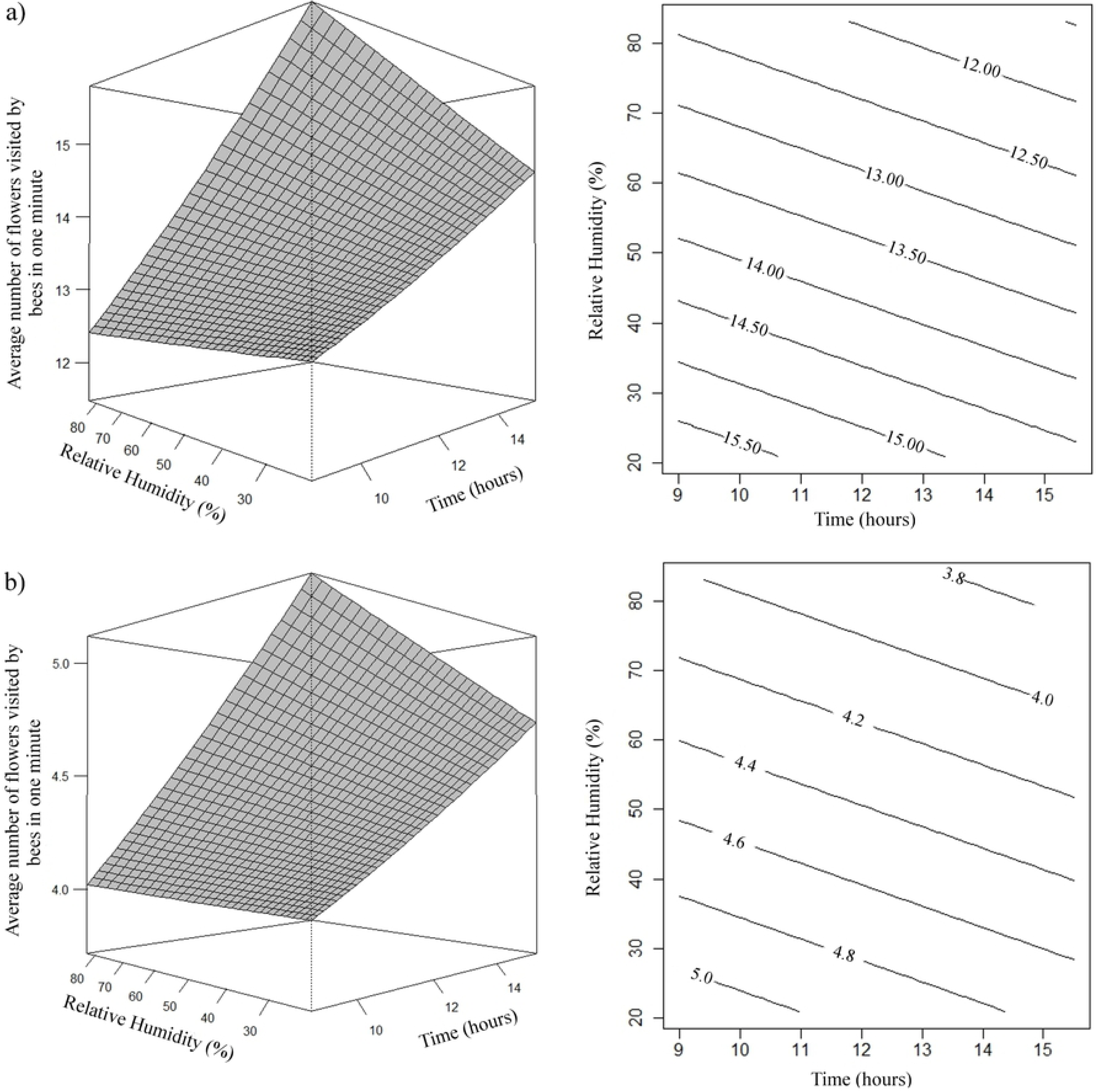
Response surface (left side) and contour (right side) graphics. Effects of daytime (x) and relative humidity (rh) on the average number of flowers visited by *A. mellifera* (a) and *T. angustula* (b) bees in one minute. (a) µ= e^2,9502-0,0120x-0,0039rh^; (b) µ= e^1,8229-0,0120x-0,0039rh^.

Estimates of the number of *A. mellifera* and *T. angustula* bees that came into contact with anthers and stigma and contact with anthers, was higher as the relative humidity increased (Fig 11 and 12, respectively). The estimated average peak of Africanized honeybees that came in contact with anthers and stigma occurred when relative humidity was approximately 76%. The highest estimates of honeybees that came into contact with the anthers occurred when the relative humidity was approximately 78%.

**Fig 11.**
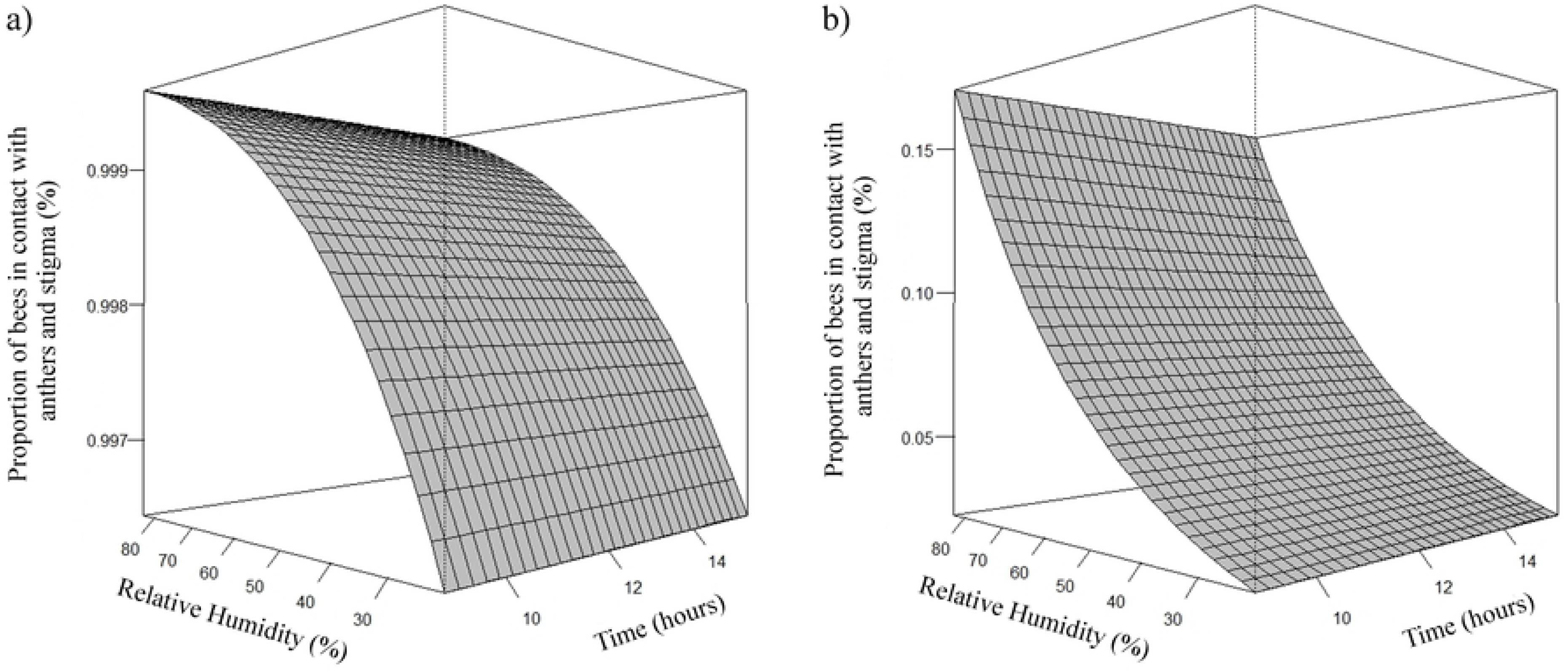
Response surface graphic. Effect of relative humidity (rh) on the proportion (μ) of *A. mellifera* (a) and *T. angustula* (b) bees in contact with anthers and stigma. (a) µ= e^4,9006+0,0350rh^/1 + e^4,9006+0,0350rh^; (b) µ= e^−4,4860+0,0350rh^/1 + e^−4,4860+0,0350rh^.

**Fig 12.**
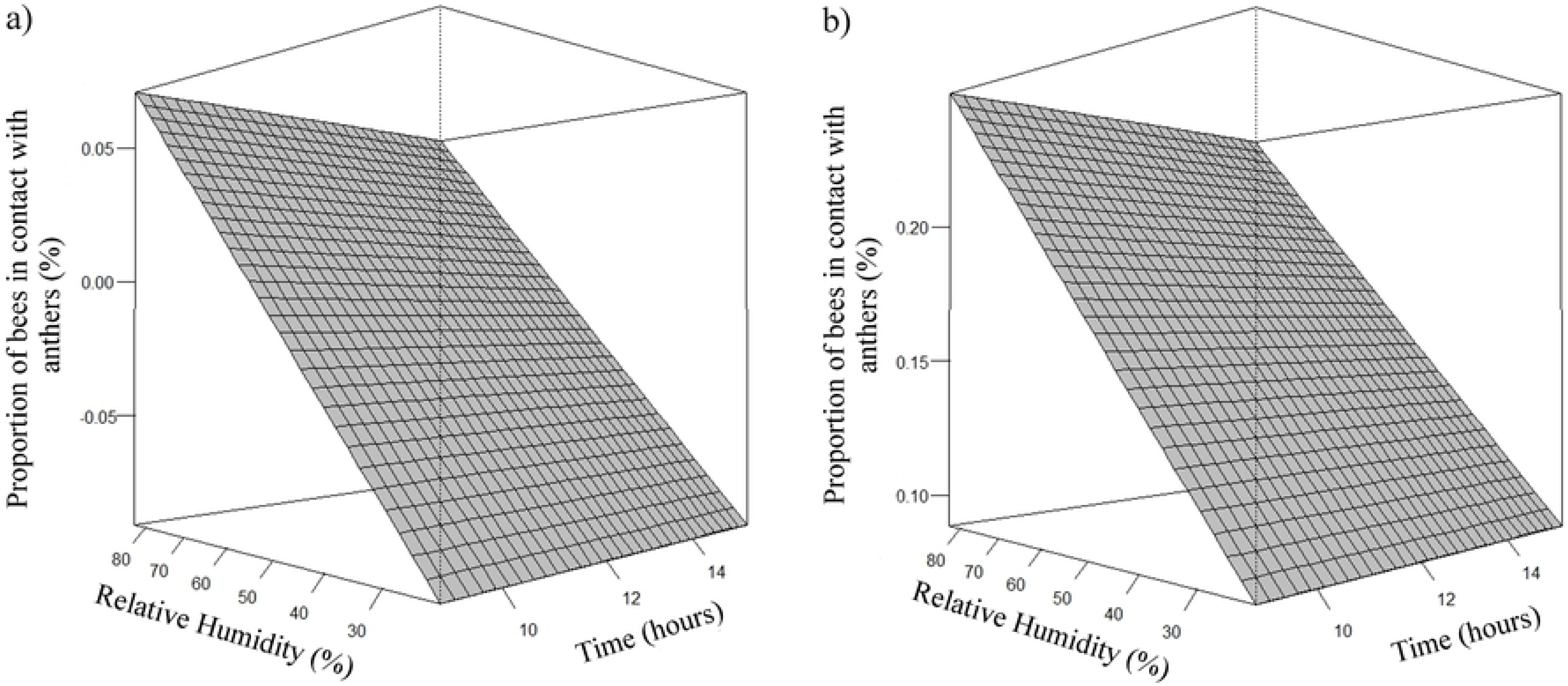
Response surface graphic. Effect of relative humidity (rh) on the proportion (μ) of *A. mellifera* (a) and *T. angustula* (b) bees in contact with the anthers. (a) µ= -0,1450 + 0,0026rh; (b) µ= 0,0341 + 0,0026rh.

## Discussion

During the foraging of canola flowers, nectar was the most sought floral resource by *A. mellifera* and *T. angustula*. Less frequent were the exclusive collection of pollen or both resources. Results similar to the behavior of *A. mellifera* were verified by [5] in *B. napus* crop and other oleaginous such as *Glycine max* [19] and *Helianthus annuus* [20]. The behavior of *T. angustula* bees was in accordance with those observed by [21] in *Citrullus lanatus* (Thunb.).

The largest number of bees collecting nectar is associated with the energy needs of the colony, since bees get all the energy necessary for their vital activities from the process of metabolizing the sugars present in the nectar. The colonies undergo large variations in the supply of nectar and weather during the year and, thus, bees collect as much nectar as possible when their supply increases, storing it in the combs in the form of honey [22].

In contrast, the supply of pollen can vary dramatically from one day to the next depending on the weather conditions and quantities of flowers, while the demand for pollen by the colony changes gradually as the number of offspring increases or decreases [22]. Therefore, the bees store what is necessary for the maintenance of the young, and, according to the colony’s need, the workers change their behavior to collect pollen, decreasing or increasing the number of foragers, size of the pollen load, and, consequently, the area of stored pollen [23].

As *T. angustula* collect nectar from the median nectaries, by the lateral base of the short stamens, they do not come in contact with the reproductive organs, considered as “stealing of nectar” since they do not pollinate the flower. This behavior was also observed in *Apis florea* species in the *B. napus* crop by [8]. However, in this experiment *T. angustula* bees were observed collecting pollen and, therefore, participating more actively in the pollination of the crop.

The number of flowers visited by *A. mellifera* honeybees and the time spent resemble those observed by [15] in *B. napus*, cultivars Hyola 61 and 433. [24] concluded that the most efficient species in the transfer of pollen grains are those that remain longer foraging the flower.

In this way, it can be suggested that the two species can increase the pollination efficiency in canola, because while *A. mellifera* is faster and has more visits, besides possessing more individuals per colony, *T. angustula* remains longer in the flower during their foraging. Because of the size of its body and its branched hairs, during foraging in the canola flowers, whether it is for collection of pollen or nectar, contact occurs with the anthers and stigmas and, consequently, cross-pollination. Cross-pollination may increase the market value of fruits and seeds by increasing quantity and quality [25].

It was observed that both species of bees preferred to collect more concentrated nectar. Nectar has a higher sugar concentration as the relative humidity decreases [26], that is, the low relative humidity provides greater evaporation of water from the nectar. Possibly, bees prefer more concentrated nectar because to turn it into honey they spend less time and have less work to dehydrate it.

The position of nectaries and exposure to the environment may result in variation in the secretion and concentration of sugars [10]. Median nectaries are more exposed, produce less nectar and are located at the base of the long stamens, while lateral nectaries are located at the base of the short stamens, in the inner part, and secrete almost all the nectar produced by the canola flower [27; 9].

The nectar produced mainly by the nectaries that are most exposed tends to reach a concentration in equilibrium with the relative humidity of the air; that is, high relative humidity further dilutes the nectars so that, consequently, there is an increase in its volume [26]. It can be deduced that as median nectaries increase the amount of nectar, bees tend to look for them more frequently. [28] reported that the nectar of lateral nectaries have higher proportions of glucose and fructose, compared to that produced in the median and as the relative humidity decreases this nectar becomes more concentrated, becoming more attractive to the bees.

With increasing temperature, the number of bees nectar collecting tends to increase for both *A. mellifera* [15] and *T. angustula* [29]. The higher number of bees in the field, greater will be the competition and extraction of nectar from nectaries. Thus, when the temperature is lower, the number of bees foraging nectar is also lower. In addition, there is a lack of nectar and pollen, increasing collection in lateral nectaries in which glucose and fructose levels are higher [28]. However, as the temperature rises, competition of foraging bees increases, reducing the availability of resources and, consequently, increasing the collection in the median nectaries.

*B. napus* flowers show higher nectar secretion rates early in the morning [27]. Environmental parameters are also responsible for the amount of nectar found during the day. These factors may have contributed to the increase in nectar collection in the median and lateral nectaries during the same visit in the afternoon, because the amount of that resource may have decreased by the decrease of secretion by the plant and due the decrease of relative humidity, forcing the bees to explore the whole flower. Availability of the nectar in the flower can present a spatial pattern, influencing the foraging movements of the pollinators on the flowers [1].

[10] observed that the maximum production of pollen occurs in the middle of the morning and early afternoon, with higher release of pollen grains by the anthers at higher temperatures, coinciding with the behavior of Africanized honeybees and *T. angustula* for pollen collection and transport observed in this research.

The time spent by the bees on the flowers will depend on the amount of floral resources offered to the pollinators, which in turn are affected by environmental parameters [26]. It has been observed that *A. mellifera* honeybees take longer collecting nectar and pollen than when collecting only nectar. Possibly, the greater time of spent on flowers during the morning period, may have occurred because there are greater floral resources available and pollen collection behavior, which was more frequent in this period.

To ensure adequate pollination and significantly increasing canola productivity rates, it is necessary to have consecutive visits to the flowers [30]. In general, plants can adjust their resources to ensure that pollinators move between as many flowers as possible [1]. Considering that the highest number of bees collecting pollen was in the morning period and that during this period there were more flowers being visited by both species studied, these characteristics showed the pollinator potential of *A. mellifera* and *T. angustula* for the canola crop.

With the increase of relative humidity, there was an increase in the number of *A. mellifera* and *T. angustula* bees that came in contact with anthers and stigma, and only with the anthers. [31] verified that *A. mellifera* honeybees actively perform pollen collection when relative humidity was between 45.00% and 89.50%. For *T. angustula*, the optimal relative humidity for external work is between 30 and 70% [29].

Africanized honeybees touched anthers and stigma during collection of nectar and pollen, while *T. angustula* only had contact with anthers and stigma during collection of pollen. During foraging there are two types of pollen collection: passive, when the pollen grains adhere to the surface of the visitor’s body; and active, when pollinators collect pollen directly from the anthers. In this case, the pollen grains used in the pollination process were not those stored in the pollen basket, but those that remain on the body of the insect [32]. During visits to the flowers pollen grains detach themselves and fall on the stigma, favoring cross-pollination; that is, *T. angustula* bees participate in pollination by the active collection of pollen.

Foraging behavior of *A. mellifera* in *B. napus* crop favored its pollination, indifferent of collected floral resource as they came into contact with anthers and stigma. *T. angustula* bees performed pollination only during pollen collection. Pollination of *B. napus* was more effective during warmer hours of the morning, when more of both species of bees carried out pollen collection. Due to its foraging behavior *A. mellifera* presented greater efficiency for pollination of *B. napus*; however, the association with T*. angustula* may potentiate the benefits generated for the crop by cross-pollination.

